# A Highly Potent SARS-CoV-2 Blocking Lectin Protein

**DOI:** 10.1101/2021.07.22.453309

**Authors:** Recep Erdem Ahan, Alireza Hanifehnezhad, Ebru Şahin Kehribar, Tuba Cigdem Oguzoglu, Katalin Földes, Cemile Elif Özçelik, Nazlican Filazi, Sıdıka Öztop, Sevgen Önder, Eray Ulaş Bozkurt, Koray Ergünay, Aykut Özkul, Urartu Özgür Şafak Şeker

## Abstract

COVID-19 pandemic effected more than 180 million people around the globe causing more than four million deaths as of July 2021. Sars-CoV-2, the new coronavirus, has been identified as the primary cause of the infection. The number of vaccinated people is increasing however prophylactic drugs are highly demanded to ensure a secure social contact. There have been a number of drug molecules repurposed to fight against Sars-CoV-2, however the proofs for the effectiveness of these drug candidates is limited. Here we demonstrated griffithsin (GRFT), a lectin protein, to block the entry of the Sars-CoV2 into the Vero6 cell lines and IFNAR-/-mouse models by attaching to spike protein of the Sars-CoV-2. Given the current mutation frequency of the Sars-CoV-2 we believe that GRFT protein-based drugs will have a high impact in preventing the transmission both on Wuhan strain as well as any other emerging variants including delta variant causing high speed spread of COVID-19.

## INTRODUCTION

SARS-CoV-2 (severe acute respiratory syndrome coronavirus 2) is the causative agent of COVID-19 (coronavirus disease-19) which has become a pandemic and global health threat since its emergence in December 2019. ^1-4^ Air-borne human-to-human transmission of SARS-CoV-2 caused the spread of the virus to almost all countries in less than a year.^5^ Due to high transmission rate, mandatory face mask use and social distancing are implemented in many countries meanwhile massive nucleic acid testing is used to suppress the further spread via finding and isolating asymptomatic and pre-symptomatic patients.^6^

SARS-CoV-2 genome is encoded in a positive-single stranded RNA molecule that has six protein coding frames consisting of ORF1ab, Spike glycoprotein (S), Nucleocapsid (N), Membrane (M) and envelope (E) proteins.^7^ S protein, which shares moderate similarity with SARS-CoV, contains a receptor binding domain (RBD) that binds ACE2 protein as the cell surface receptor to initiate viral invasion.^8, 9^ S protein is found as a trimer with two metastable conformations in which RBD is either “up” or “down” state. Upon binding to ACE2, S1 and S2 domains of S are cleaved by surface protease such as TMPRSS2 or cathepsin L. Cleavage of S protein domains causes irreversible structural changes and primes for the viral fusion.^10-12^ Due to high antigenicity of S proteins and importance of ACE2 binding for virus entrance, many prophylactic vaccines based on mRNA (BioNTech/Pfizer^13^, Moderna^14^), adenoviral vector (J&J^15^, AstraZeneca/Oxford^16^) and recombinant antigen (Novavax^17^) utilize S protein to induce anti-SARS-CoV-2 immune response.

Aside from vaccines, current treatment options of COVID-19 include disruption of viral amplification by either inhibiting viral entrance to cells or viral replication machinery. Drug repurposing studies successfully identified several small molecules such as remdesivir as a viral replication inhibitor that acts on RNA polymerase of SARS-CoV-2.^18^ Moreover, neutralization antibody (nAbs) therapies covering convalescent plasma transfer and recombinant monoclonal anti-Spike antibodies are also approved for emergency use. ^19, 20^ Generally, nAbs render the binding of RBD to ACE2 protein thereby preventing the virus internalization. Yet, some nAbs bind regions other than RBD in S protein to hinder conformational changes for fusion state.^21^ Although apparent beneficial outcomes of nAb therapies among COVID-19 patients, limited availability of convalescent plasma and costly production process of monoclonal antibodies restrain accession to this treatment.^22^ In addition, ongoing evolution of SARS-CoV-2 poses a significant risk as mutation accumulation in S protein can lead to immune escape which can cause reinfections and make nAbs therapies as well as vaccination ineffective.

Recent reports showed that three variant of concern (VOC) strains which are B.1.1.7 (England), B.1.351 (South Africa), and P.1 (Brazil) circulating in the globe have moderately enhanced transmission and/or immune evasion characteristics from ancestral strain, D614G through acquired mutations such as E484K and N501Y.^23^ Full evolution potential of S protein is still needed to be determined, yet it is highly-likely that SARS-CoV-2 will acquire more mutations to cope with pressure due to vaccine-elicit immunity because uneven distribution of vaccines provide viable hosts for evolution.^24^ Hence, new therapeutic agents that have distinct inhibitory action are required until complete eradication of SARS-CoV-2.

Lectin proteins isolated from seaweeds are shown to be potent antiviral agents against enveloped viruses e.g., HIV-1^25^, herpes virus^26^ as well as two deadly human coronaviruses, SARS-CoV^27^ and MERS-CoV^28^. Antiviral activity of seaweed lectins arises from their affinity to surface glycoproteins on viruses such as gp-120 protein of HIV-1 and spike proteins of SARS-CoV and MERS-CoV. Upon binding to surface proteins, lectins generally block the viral internalization step and thereby prevent the viral infection. In the present study, we evaluated the activity of griffithsin lectin protein (GRFT) from *Griffithsia sp*. against the novel human coronavirus, SARS-CoV-2. For this purpose, GRFT is recombinantly expressed in *E. coli* with histidine tag and purified. Binding of recombinant GRFT to whole inactivated SARS-CoV-2 as well as purified spike protein from HEK293 are validated and characterized with ELISA, ITC and QCM. The activity of GRFT is assessed *in vitro* with Vero6 cells, and *in vivo* with Syrian hamsters. Our results indicate that GRFT is a potent non-mutagenic antiviral agent against SARS-CoV-2, reducing virus transmission through blocking its entry into the cells. Although the vaccination of the world population is in progress, the time to reach herd immunity is still unknown. Also, recent mutation reports on different subunits of the SARS-CoV-2 may urge the need for changing the current vaccine designs. In this regard our proposed antiviral GRFT can help to suppress the transmission of the virus. Prevention of person-to-person transmission may also help to stop the evolutionary change of the virus through selective pressure. Upon very promising results from *in vitro* and *in vivo* assays and experiments, GRFT is formulated as a nasal spray for upcoming human phase trials. We believe that GRFT based nasal spray will have the potential to change the current scenario of the pandemic.

## RESULTS

Spike glycoprotein is heavily glycosylated with oligomannose and complex type sugars on its estimated 22 N-linked and 2 O-linked potential glycosylation sites per protomer. Total glycans structures, when recombinantly expressed in HEK293, account for one-to-third molecular weight of Spike protein and cover approximately 40% of the protein surface. ^29, 30^ (Figure 1a) Owing to high conservation of sequons among SARS-CoV and SARS-CoV-2 spike proteins^31^, we hypothesized that GRFT protein can act as an antiviral for SARS-CoV-2, due to its previously reported antiviral effects on SARS-CoV. GRFT is considered as a domain swapped dimer folded as beta prism which can bind 3 sugar molecules per protomer.^32^ (Figure 1b)

**Figure 1.**
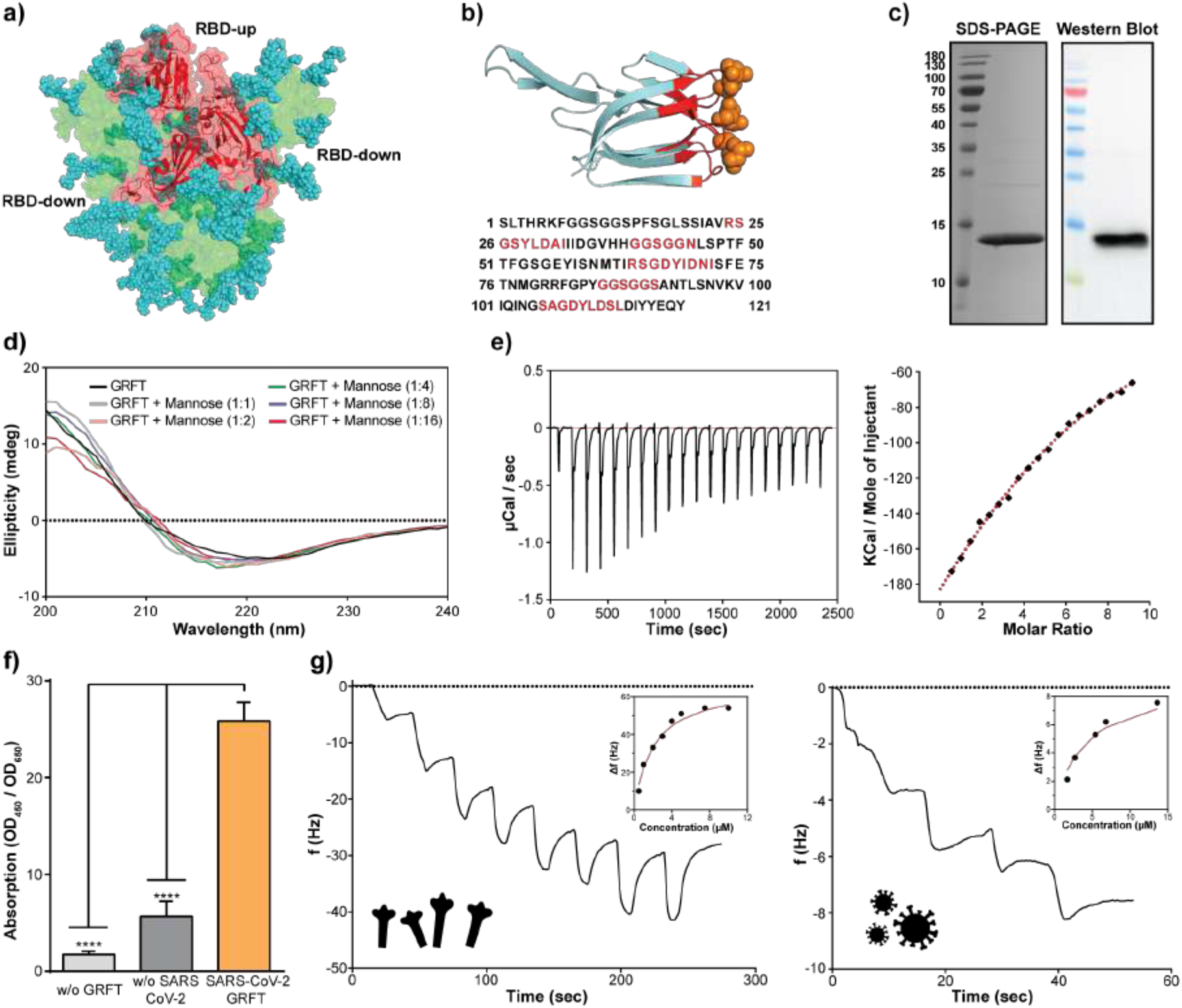
**a**. Structure of HEK293 produced recombinant SARS-CoV-2 spike protein, and receptor binding domain with attached glycan structures colored as blue (side view)^30^ **b**. Structure and amino acid sequence of rGRFT **c**. Purification validation of rGRFT protein on SDS-PAGE and Western Blotting gels, single bands are representing the expected 14.5 kDA molecular mass of GRFT **d**. Secondary structure changes of rGRFT upon titration with mannose **e**. the molecular binding interaction of SARS-CoV-2 spike protein with rGRFT protein analyzed with ITC f. Qualitative ELISA assay for the interaction of rGRFT protein with SARS-CoV-2 virus g. QCM-D based quantitative analysis of the binding of rGRFT on purified spike (left) and inactivated SARS-CoV-2 (right) immobilized on carboxylated gold surface

To test our hypothesis, recombinant GRFT (rGRFT) was expressed with 6x His-tag in *E. coli* BL21 (DE3) and was purified with Ni-NTA column. Final yield of pure rGRFT was found approximately 12mg/L consistent with previous reports at shake-flask scale in LB media.^33^ Purity of rGRFT was calculated as >95% via SDS-PAGE under non-reducing conditions. rGRFT purification was further validated with Western Blot using antibody against his-tag. (Figure 1c) The secondary structures of rGRFT were measured with circular dichroism both in apo-form and complex with mannose sugar. In the CD spectrum, apo-rGRFT gave minima around 218nm in agreement with its beta-sheet rich 3D structure. Overall secondary structure elements were not changed upon addition of mannose up to 16-fold molar excess over rGRFT (Figure 1d, Supplementary Table 1). Binding of rGRFT to recombinant SARS-CoV-2 spike was investigated with isothermal calorimetry (ITC) and quartz crystal microbalance (QCM) techniques. Dissociation constant (K_d_) of rGRFT-Spike protein complex was calculated as 9.9μM and 1.6μM from ITC and QCM-D, respectively. ITC results provided that binding stoichiometry between rGRFT and Spike is 6:1 (Figure 1e and Figure 1g-left panel). Next, binding of rGRFT to the whole virus was analyzed with in-house developed ELISA in which heat-inactivated SARS-CoV-2 particles were coated onto wells to capture free rGRFT. In presence of SARS-CoV-2 particles, rGRFT remained bound to wells after washing, meanwhile statistically less rGRFT was detected onto wells in absence of SARS-CoV-2 particles. (Figure 1f). QCM-D was utilized to determine the binding affinity of rGRFT to the viral particles. rGRFT can bind heat-inactivated SARS-CoV-2 viral particles with K_d_ of 4.1μM (Figure 1g-right panel).

Efficacy of rGRFT against SARS-CoV-2 infection was assessed *in vitro* by using the VeroE6 model cell line. (Figure 2a) Serial diluted rGRFT solutions at concentrations ranging from 6μM to 15.4pM were mixed with infectious SARS-CoV-2 particles. Subsequently, cells were infected with mixtures and incubated until the cytopathic effect (CPE) reading reached %100 in the sample that didn’t include rGRFT. Based on the infection results, EC50 of rGRFT on SARS-CoV-2 infection was found 5.76 nM. (Figure 2b) Effect of rGRFT at EC50 on SARS-CoV-2 replication was analyzed as well. VeroE6 cells were infected with SARS-CoV-2 in presence and absence of rGRFT and the amount of total viral particles in cell supernatant was determined using qPCR. rGRFT is able to suppress SARS-CoV-2 infection after at most 24 hours upon inoculation (Figure 2c). In addition, rGRFT did not show any cytotoxicity effect on cells at concentration as high as 137nM. (Figure 2d)

**Figure 2.**
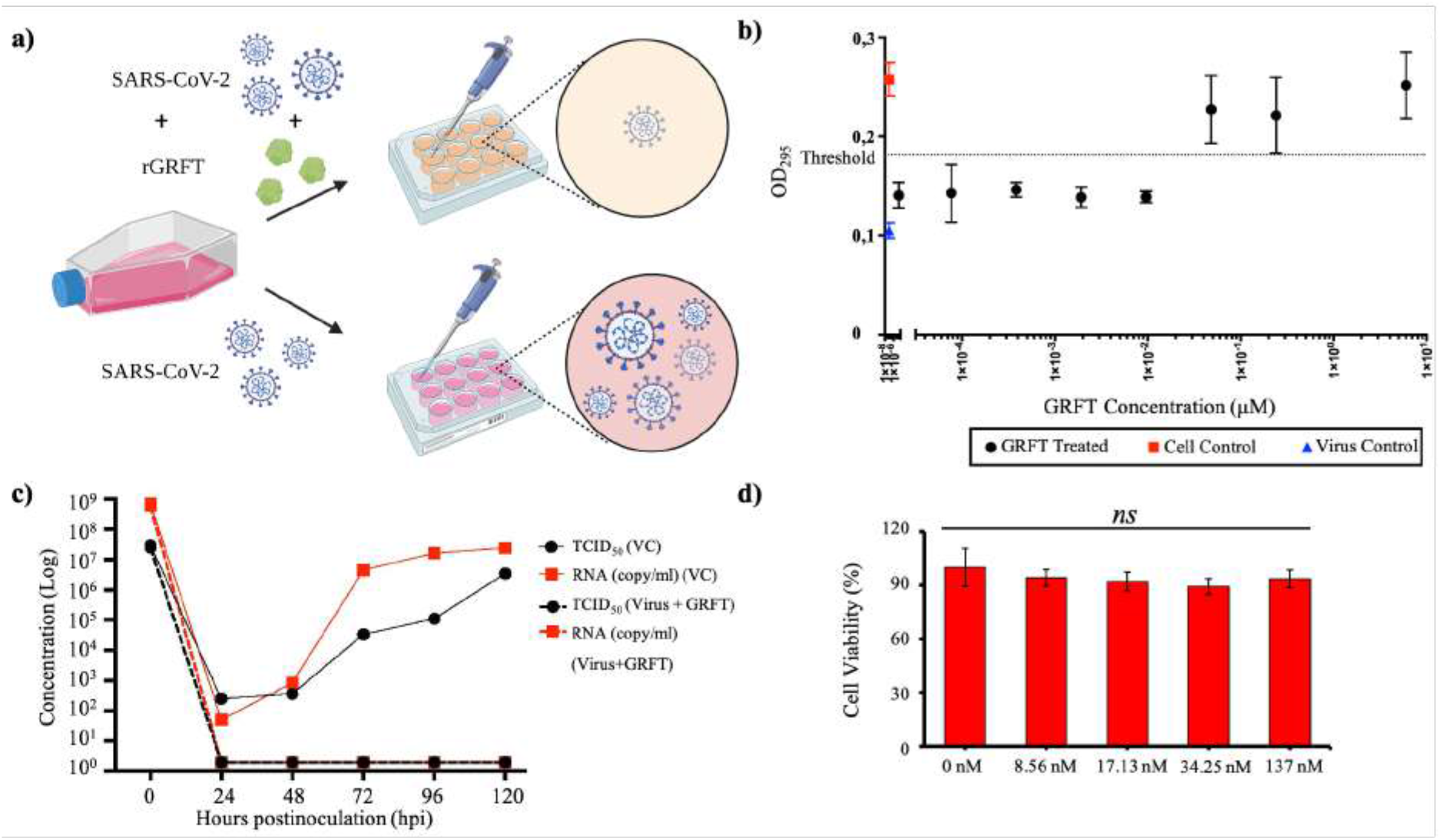
**a**. Schematic representation of rGRFT inhibiting SARS-CoV-2 infection in Vero E6 cells. Created with BioRender.com. **b**. Determination of rGRFT EC50 value for SARS-CoV-2 in Vero E6 cells. **c**. Effects of rGRFT (5.76 nM) on SARS-CoV-2 replication in Vero E6 cells. d. Cytotoxicity assessment of rGRFT on Vero E6 cells by MTT assay.

Prior to efficacy experiments *in vivo*, toxicity and immunogenicity of rGRFT on live C57BL/6 mice was investigated. 150μl of 100nM rGRFT was injected to mice intraperitoneally and the mice were monitored for 14 days. rGRFT injection didn’t cause any meaningful variation in body temperature and weight of mice. (Figure 3a and Figure 3b). During the 14-day observation period, mice did not show any evidence for local or systemic toxicity. Furthermore, biochemical parameters for liver and kidney and blood count results remained within the healthy range. (Table 1 and Table 2). Tissue samples of liver, lung, kidney, and spleen obtained from sacrificed animals were examined under microscope to assess histopathological effects. No visible histopathological changes were observed from corresponding tissue samples. (Figure 3d) Immunogenicity of rGRFT was analyzed with in-house developed ELISA that capture anti-rGRFT antibodies from serum via surface immobilized rGRFT. ELISA results showed that rGRFT did not simulate the humoral immune response. (Figure 3c).

**Figure 3.**
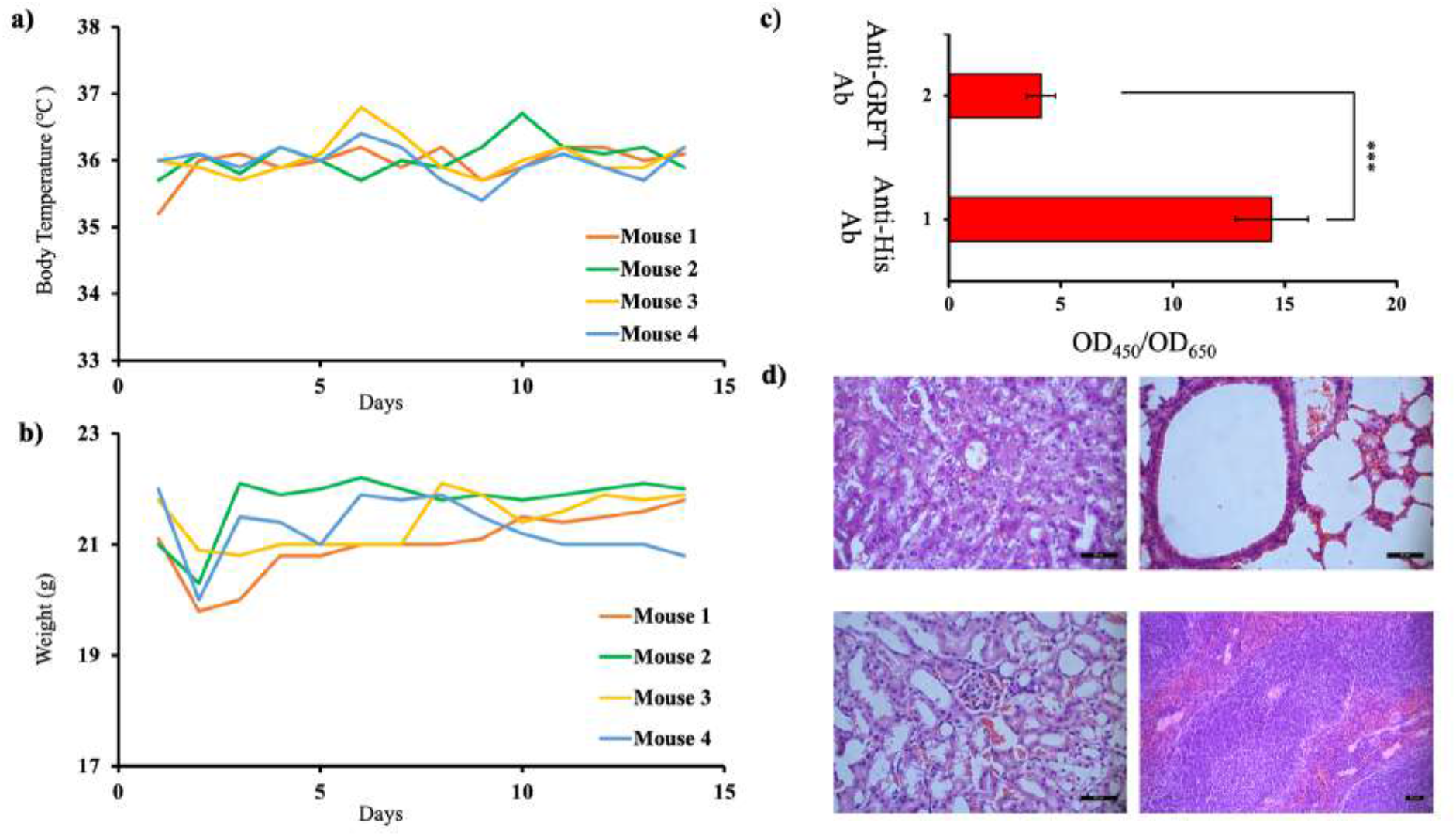
**a**. Body temperature and **b**. weight recorded for 14 days in C57BL/6 mice during innocuity testing. **c**. Immunogenicity of rGRFT administered to C57BL/6 mice during innocuity testing. Antibodies against rGRFT were significantly lower than those induced by the His-tag in the construct (p<0.05). **d**. Representative tissue sections of liver (upper left), lung (upper right), kidney (lower left) and spleen (lower right) in C57BL/6 mice on 14th day following intraperitoneal rGRFT administration (hematoxylin and eosin staining, scale bar: 50 µm).

*In vivo* efficacy of rGRFT was tested in two different experimental set-ups. In the first set-up, rGRFT was administered intranasally to mice prior to direct virus inoculation through the nose. In the second set-up, mice were separated into three groups; the first mice group were administered with rGRFT, the second mice group remained untreated, and the third mice group were infected intranasally with virus particles. (Figure 4a) Subsequently, each member of the third mice group was put together into a cage with either a member of the first mice group or a member of the second mice group. Treatments of each individual mouse were repeated for 6 days. Mice were monitored for 14 days to assess the protective role of rGRFT. In both scenarios, admission of GRFT decreases the viable infectious particle number as well as viral load determined with qPCR. (Figure 4b)

**Figure 4.**
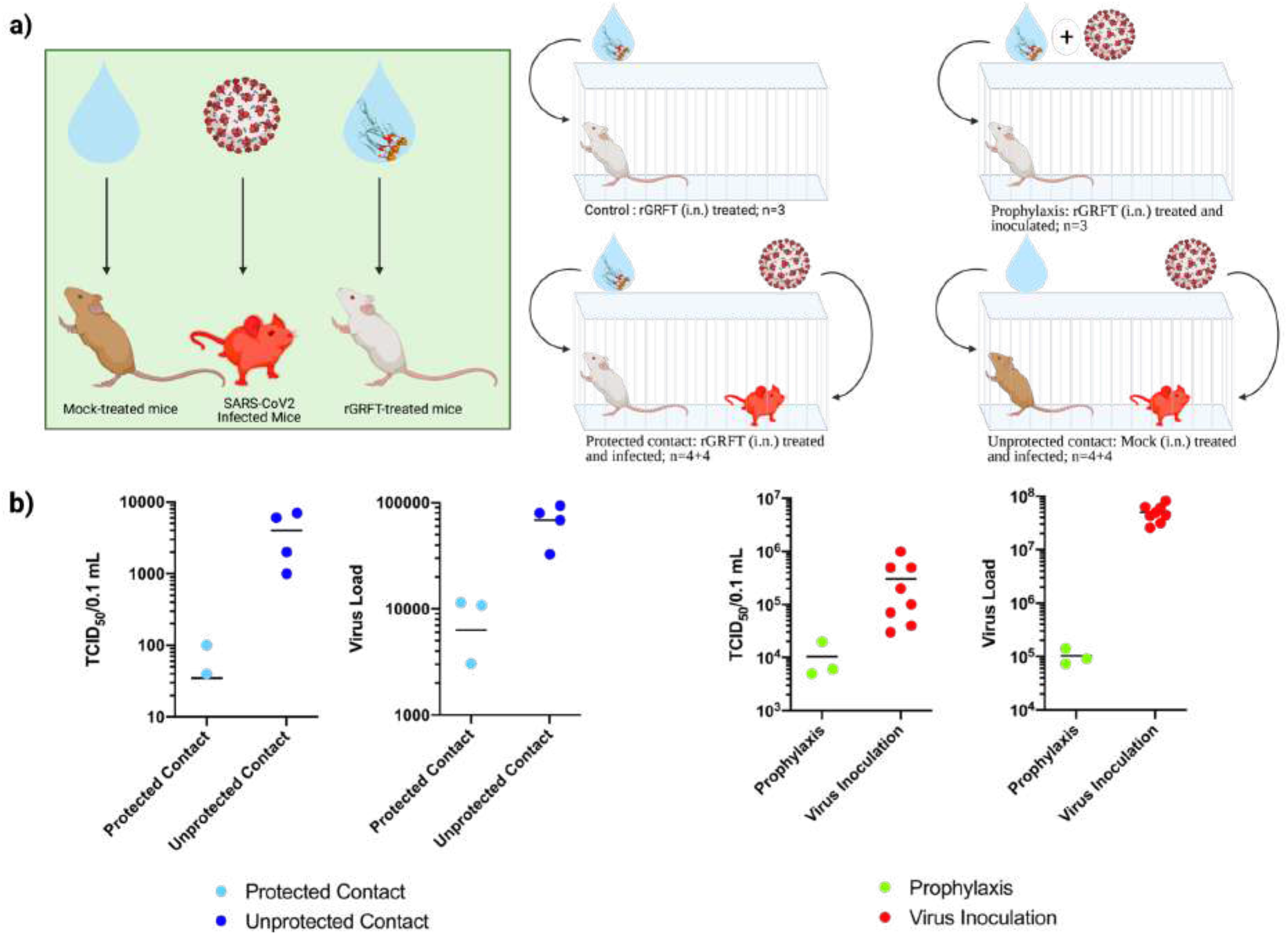
**a**. IFNAR-/-mice assigned to various study groups. Control group is mice treated with only rGRFT (i.n.) in a cage (n=3), prophylaxis group is rGRFT (i.n.) treated and SARS-CoV-2 inoculated mice in a cage (white mice; n=3), protected contact group is consisted of rGRFT treated mice and SARS-CoV-2 infected mice (white mice and red mice, respectively; n=4+4), unprotected contact group is mock (i.n.) treated mice and SARS-CoV-2 infected mice (brown mice and red mice, respectively; n=4+4). **b**. SARS-CoV-2 loads and infective virus titres in lung tissues of individual IFNAR-/-mice assigned to various study groups. rGRFT treated mice from protected contact group, mock treated mice from unprotected contact group are compared in order to SARS-CoV-2 loads and infective virus titres; prophylaxis group (rGRFT treated and SARS-CoV-2 infected mice) and all SARS-CoV-2 infected mouse are compared in order to SARS-CoV-2 loads and infective virus titres. Protected contact group values are represented with red color; unprotected contact group values are represented with blue color; prophylaxis group values are represented with green color; virus inoculation group is represented in orange color.

On day of experiment termination, SARS-CoV-2 infection was monitored in lung tissues by in situ hybridization test (ISH). The genomic RNA of the virus was detected more diffuse in lungs of unprotected contact animals when compared to those protected Pattern of the infection in protected contact animals appeared patchy. SARS-CoV-2 RNA was detected in bronchial epithelial cells, alveolar epithelial cells type I and type II, as well as macrophages in both groups. (Figure 5).

**Figure 5.**
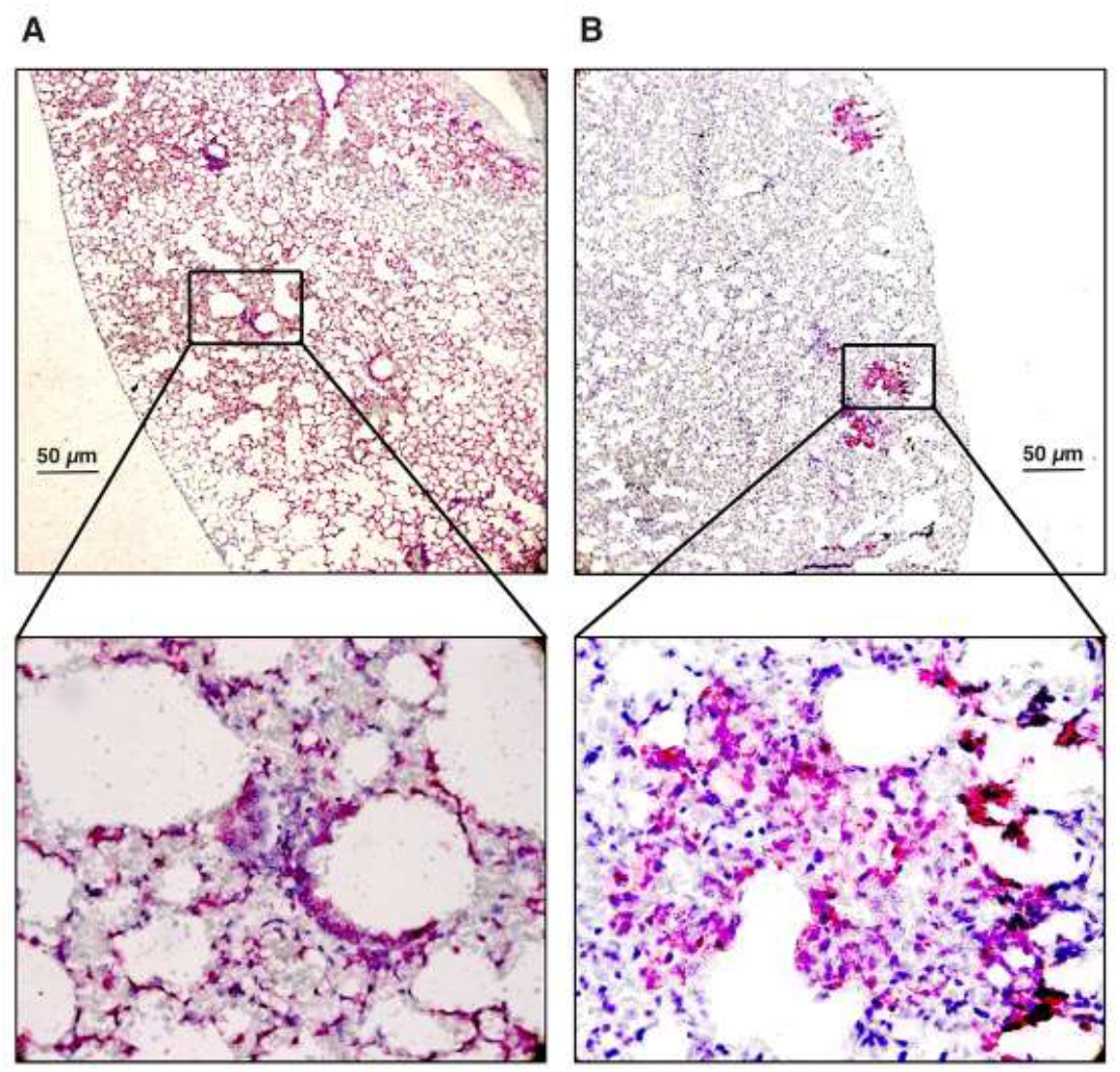
In situ hybridization of viral RNA in unprotected (A) and protected (B) animals

## DISCUSSION

COVID-19 pandemic is an ongoing concern for all countries that put a significant burden on healthcare systems and the economy. Owing to endless efforts of researchers, governments and pharmaceutical companies, viable vaccine options against the SARS-CoV-2 are available.^34-39^ Alternatively, neutralization antibody therapies along with antibody mimetic peptides raise a hope to end the current pandemic in the foreseeable future.^19, 40-43^ Yet, continuous evolution of SARS-CoV-2, especially mutations occurring in the spike protein region, pose a risk to lose or minimize the efficacy of both vaccines and nAbs. Currently, several virus strains such as P.1 and B.1.351 can evade neutralization antibodies to a certain degree which are determined with *in vitro* pseudovirus assays.^44^ Despite, the evolution capacity of SARS-CoV-2 S protein is unknown hitherto, low homology with other ACE2 recognizing coronaviruses such as HCoV-NL63 implied that S protein can adapt different mutations without loss-of-function^24^. This worrisome scenario necessitates broad ‘pan-coronavirus’ inhibitory agents to suppress the transmission until a viable vaccine is available for emerging variants.

Here, we demonstrated that griffithsin protein from *Griffithsia sp*. recombinantly produced in *E. coli* can bind S protein of SARS-CoV-2 *in vitro* and inhibit its infection both *in vitro* VeroE6 cell line and in *vivo* mouse model when applied prophylactically. Toxicity assays of rGRFT with mouse models indicated that it is a tolerable agent even at concentrations higher than its therapeutic concentration window.

Previous studies showed that rGRFT can inhibit both MERS and SARS infection *in vitro* cell models.^27, 28^ The S protein of SARS-CoV-2 shares low-to-moderate similarity with clinically relevant MERS-CoV and SARS-CoV ^45^, yet it is proposed that the inhibitory mechanism is based on binding to glycan structures rather than amino acids sequence motifs.^27^ These glycan structures, in contrast to general amino acid sequence, are conserved between SARS-CoV and SARS-CoV-2. In addition, glycan sequons of SARS-CoV-2 are reported to be conserved in the course of Covid-19 pandemic between December 2019 to April 2020.^46^ This phenomenon is correlated with another report which showed that SARS-CoV-2 mutants without glycans especially at N165 and N234 sequons have decreased affinity to ACE2 protein. In the report, authors claimed that loss of glycans at stated sites shifts the equilibrium of RBD states to down position which might lead to evolution of less infectious particles.^47^ Therefore, there might be natural pressure on SARS-CoV-2 for conservation of N-glycan which can be exploited via rGRFT treatment to hinder viral transmission.

Aside from efficacy, rGRFT production cost is reported to be as low as 3,500$ per kg with yield approximately 2,5 g/L in *E. coli* at scale of tons.^48^ In addition, rGRFT is reported to be stable at room temperature in PBS for 2 years without formation of aggregation and degradation byproducts.^49^ Considering the board activity of rGRFT on different coronaviruses, evolutionary cost of glycan loss on SARS-CoV-2, available high-yield recombinant production strategies for rGRFT and good stability of rGRFT at room temperature, rGRFT may provide a possible solution for emerging variants.

## METHODS

### Cloning, Expression and Purification of Griffithsin Protein

The griffithsin (GRFT) nucleotide sequence (accession number AY744144) with N-terminal 6X his tag and flexible GS linker was codon optimized for *Escherichia coli (E. coli)* and chemically synthesized (Genewiz, NJ, USA) with suitable overhang sequences homologous to pet22b cloning vector, without the pelB leader sequence. The GRFT amino acid sequence was listed in Supplementary Table 4. *grft* gene was cloned into the pet22b vector without a pelb leader sequence, using Gibson assembly. The reaction was transformed into a chemically competent strain of *E. coli DH5α PRO* by heat shock transformation. Selected colonies were verified by Sanger sequencing (Genewiz, USA).

For expression, the construct was transformed to *E. Coli BL21 (DE3)* strain. Overnight culture of cells was diluted to 1:100 in LB medium with appropriate antibiotics and was grown till OD_600_ was 0.4-0.6. Then, the culture was induced with 1 mM IPTG for 24 hours at 16 °C, 200 rpm. Cells were harvested by centrifugation and resuspended in the lysis buffer (20 mM NaH2PO4, 500 mM NaCl, 20 mM imidazole pH 7.4). The suspension was sonicated at 30% power for 10 cycles of 15s on/45s off. Samples were centrifuged at 21.500g for 1 hour and supernatant was filtered with 0.45μm filter. Filtered lysate was loaded on a pre-equilibrated HisTrap nickel column (GE life sciences 17524701) using ÄKTA Start protein purification system. The column was washed with 10 column volumes of the lysis buffer (20 mM imidazole) and 5 column volumes of the lysis buffer with an imidazole gradient from 20 mM to 40 mM. Finally, proteins were eluted with 5 volumes of elution buffer (20 mM NaH2PO4, 500 mM NaCl, 500 mM imidazole pH 7.4). Purified Griffithsin was desalted into 1 X PBS (pH 7.4) using HiTrap desalting column (GE Lifesciences) in ÄKTA Start protein purification system.Protein concentration is calculated via BCA colorimetric assay with BSA standards. (ThermoFisher Scientific)

### SDS-PAGE and Western Blotting

Samples were boiled at 95 °C for 5 minutes with 1 X SDS loading dye and electrophoresed on 15% SDS-polyacrylamide gel. The gel was placed into the Coomassie blue staining solution for ∼1 hour with shaking and incubated in destaining solution (60 % ddH_2_O, 30% methanol, 10% acetic acid) until the bands were clearly visible. For Western blot, gel was transferred to the PVDF membrane using Trans Blot Turbo (Biorad). The membrane was blocked with 5% milk in TBS-T for 1 hour at room temperature. Then the membrane was incubated in 5% milk in TBS-T containing 1:10000 primary anti-his mouse antibody at 4°C, overnight. Membrane was washed in TBS-T, then incubated in 5% milk in TBS-T containing 1:10000 horseradish peroxidase (HRP) conjugated goat anti-mouse secondary antibody (Abcam ab6789-1 MG) for 1 hour at room temperature. After washing in TBS-T, the membrane was incubated in ECL substrate (Biorad 170-5060) and visualized using Vilber Fusion Solo S.

### Circular Dichorism (CD)

7.5 µM of rGRFT was prepared in 2mM PBS buffer pH=7.4. Mannose concentrate solution was prepared with ddH_2_O and diluted in GRFT solution at 7.5, 15, 30, 60 and 120 µM final concentration. The mannose protein mixture was incubated for 30 minutes at room temperature. The CD spectra of GRFT/mannose mixtures was measured from 240 nm to 200 nm with 3 repeats (Jasco J-815) at room temperature with 300 sec delay time, 1 mm band width. Secondary structure composition was calculated using the BestSel tool developed by Micsonai et al.^50^

### Isothermal Titration Calorimetry (ITC) Assay

Isothermal titration calorimetry assays were performed using MicroCal ITC200 (Malvern Panalytical). Spike protein was purchased from Synbiotik Biotechnology LLC (Ankara, Turkey). 1 μM of spike protein solution and 50 μM of rGRFT protein solution were prepared in 1x PBS, pH=7.4. 2 μl of rGRFT solution was injected with 120 seconds intervals into the calorimetric cell containing 280 μl of Spike protein solution. Buffer and dilution effects were corrected using Origin MicroCal Analysis Software. Graphs were generated via GraphPad Prism software.

### Quartz Crystal Microbalance (QCM) Assay

QCM-D gold sensor (BiolinScientific QSense^®^ QSX 301 Gold) surface was cleaned by piranha solution (H_2_O_2_/H_2_SO_4_ in 1:3 ratio) for 30 minutes at 80 °C to remove any contaminants. After piranha cleaning, the gold sensor chip was immersed in ddH_2_O for 5 minutes twice.

The cleaned gold sensor chip was immersed in 20 mM 11-Mercaptoundecanoic acid (11-MUA) solution and incubated overnight. The gold sensor was rinsed with first 100% ethanol then ddH_2_O. The gold sensor surface coated with 11-MUA was functionalized by QSense chamber with 400 mM EDC which was followed by 100 mM NHS with a flow rate of 20 µL/min. The chip was rinsed with 1X PBS buffer to remove any residual EDC and NHS. Then, 150 µL Spike protein with a con°centration of 400 µg/mL was introduced to the chamber with 6.49 µl/min flow rate to make Spike protein attached to the surface. After coating the surface with Spike protein, the surface was deactivated by 1M ethanolamine HCl with a flow rate of 20 µL/min to avoid further attachment to the functionalized surface rather than Spike protein. Deactivation was followed by 1X PBS washing step to equilibrate the chamber at 20 µl/min. After, 200 µl of 0.5 µM, 1 µM, 2 µM, 3 µM, 4 µM, 5 µM, 7.5 µM, 10 µM of rGRFT was introduced sequentially at 20 µl/min flow rate. Between each concentration, chip was washed with 1X PBS at 20 µl/min flow rate to remove any unbound molecule. 1^st^, 3^rd^, 5^th^, 7^th^ and 9^th^ overtones for frequency and dissipation values were recorded simultaneously.

Binding affinity of rGRFT towards SARS-CoV-2 virus was tested with QCM-D. The surface of the gold sensors was prepared as described above. The viral particles were attached on the surface of QCM-D via amine coupling. After the attachment of the viral particles the activated surface as mentioned above blocked with ethanolamine (1M) in PBS. Final surface topographies were analyzed with atomic force microscopy as demonstrated elsewhere.^51^ Afterwards, rGRFT proteins at varying concentrations as following 1.7 µM, 2.7 µM, 5.4 µM, 6.8 µM, 13.6 µM were flown on top of the inactivated and immobilized Sars-CoV-2 virus on sensor chips.

### rGRFT Binding on SARS-CoV-2 by ELISA

96 well plates (353916, Corning) were coated with 50 µL of 10^6^ particle per mL SARS-CoV-2 isolated from VeroE6 cells by adding 150 µL 100 mM Bicarbonate/carbonate coating buffer (pH 9.6) and incubated O/N at 4°C. Wells were washed 3 times with 200 µL PBS containing 0.1% Tween-20 (Merck) (PBS-T) 200 µL blocking buffer (1% BSA in PBS-T) were added to wells and incubated for 1 hour. After incubation, the blocking buffer was discarded and 100 µL of 1µg/mL rGRFT was added to each well and incubated for 2 hours. Wells are washed with 200 µL PBS-T three times. Then, each well was incubated with a PBS-T solution containing 1:3000 primary anti-his mouse antibody (PTGLAB 66005) for 1 hour. Washing was performed with 200 µL PBS-T. Secondary antibody solution containing 1:3000 horseradish peroxidase (HRP) conjugated goat anti-mouse secondary antibody (Abcam ab6789-1 MG) in PBS-T was added to each well and incubated for 1 hour. Washing steps were performed with PBS-T and wells were incubated with 100 µL TMB substrate solution for 10 mins in dark. TMB stop solution was added and absorbance at 450 nm and 650 nm was measured with microplate reader. (SpectraMax M5, Molecular Devices).

### Cell and Animal Experiments

#### Virus, Cells, Animals and Ethical Approval

SARS-CoV-2 Ank1, a previously characterized local isolate, was used in the experiments requiring live virus. ^51^ African green monkey kidney (Vero E6, ATCC: CRL-1586) cells, obtained from the cell culture collection of the Department of Virology, Ankara University Faculty of Veterinary Medicine were also utilized in the study. The experiments with C57BL/6 and IFNAR-/-mice were performed with permission of the Ankara University Ethical Committee for Animal Experiments (06 May 2020, 20120-8-66) in the high containment animal facility (ABSL3+), conducted according to the national regulations on the operation and procedure of animal experiments’ ethics committees (regulation no. 26220, 9 September 2006). The 14-day non-lethal SARS-CoV-2 infection model in IFNAR-/-mice were previously reported. ^51^

Tissue culture infective dose 50% (TCID_50_) was used to assess in vitro virus infectivity, performed as described previously.^51, 52^ Quantitative genome detection was carried out by real-time reverse transcription polymerase chain reaction as reported.^51^

### Toxicity and immunogenicity

Vero E6 cells were grown in Dulbecco’s modified Eagle’s medium (DMEM) supplemented with 10% heat-inactivated fetal bovine serum, 2 mM L-glutamine and 100 units/ml penicillin, 100 µg/ml streptomycin, at 37 °C in a 5% CO2 humidified incubator. To determine toxicity of rGRFT on Vero E6 cells, MTT (3-[4,5-dimethyl-2-thiazolyl]-2,5-diphenyl-2H-tetrazolium bromide) assay was used. 3 ×10^5^ cells were seeded onto 96 well plates and incubated at 37 °C in a 5% CO2 humidified incubator for 24 hours. Then, different concentrations of rGRFT in PBS were added on cells, and incubated for another 48 hours. Cell viability was assessed using the MTT Cell Proliferation Assay Kit (Trevigen, 4890-25-K), according to manufacturer’s instructions..

For innocuity testing, four C57BL/6 mice were intraperitoneally inoculated with a 100nM (250 µL) rGRFT, three times with 24-hour intervals and observed for 14 days with daily measurements of body temperature and weight. They were sacrificed at the end of the period with specimens for biochemical, hematological and histopathological assessment. Mouse sera at day 14 was further screened for rGRFT-specific antibodies. For this purpose, ELISA plates were coated overnight with rGRFT in bicarbonate buffer and the sera in 1/100 dilution were inoculated for an hour at room temperature. The assay was evaluated using anti-mouse-IgG-HRP conjugate and TMB substrate. The x6-His peptide present in the recombinant protein was targeted as the control, detected using mouse anti-His antibodies, anti-mouse-IgG-HRP and conjugate and TMB substrate.

### In vitro testing of rGRFT

In vitro virus inhibition was monitored by two approaches. Initially, neutralizing activity of rGRFT at different incubation periods was determined. For this purpose, rGRFT (6 µM) was 5-fold diluted in high glucose DMEM, combined with an equal volume (100 µL) 100 TCID_50_ virus and the mixture was incubated at room temperature for 15 minutes. A 150 µL of the mixture was then inoculated on Vero E6 Cells grown in a 96-well flat-bottomed tissue culture plate with highest concentration (6 µM) of rGRFT without the virus was involved as toxicity, serum-free DMEM as cell and 75 µL 100 TCID_50_ virus as virus controls in each plate. The plates were incubated at 37°C in 5% CO_2_ atmosphere and evaluated when the virus controls showed 100% CPE. The 50% effective dose (EC_50_) was calculated as described elsewhere.^27^

In the second approach, Vero E6 cells grown in 24-well tissue culture plates were infected with SARS-CoV2 Ank1 isolate at 1 moi and rGRFT (in EC_50_ concentration) was added at 1-hour post-infection. The test was incubated until virus controls showed 90% CPE. TCID_50_ values and virus genome copies were determined in culture supernatants and cell layers at termination. The test was performed in quadruplicate for all time points.

### In vivo testing of rGRFT

rGRFT was delivered intranasally in 100 nmol per nostril in 50 µL volume of carrier solution including 2.5% (w/v) polyethylene glycol 1450 (PEG 1450), 0.5% benzyl alcohol, 1.5% (w/v) glycerine, 0.02% (w/v) h) EDTA-diNa and 0.02% (w/v) benzalkonium chloride. The solution was sterilized by membrane filtration (0.2 nm, Milipore, USA) and stored under chilled conditions. Intranasal virus inoculations were carried out as 10^3^TCID_50_ in 50µL per nostril.

A total of 28 IFNAR^-/-^ mice (8-12 weeks) were randomly divided into 6 groups for prophylaxis and contact experiments as well as controls Individuals in the prophylaxis group received intranasal rGRFT and inoculated with SARS-CoV-2 Ank1 after 4 hours. Each animal further received the same amount and route of rGRFT following 4 subsequent days. The protective effect of rGRFT was tested with a contact scenario in pairs housed in the same cage. One individual in each cage was intranasally inoculated with the virus and the accompanying mouse received rGRFT (protected contact) or null carrier solution (unprotected contact) (Figure 2). The rGRFT and carrier treatments were repeated at the same time for 6 additional days. Animals were monitored clinically for 14 days with tail vein blood specimens collected on days 0 and 7. At the end of the period, the animals were humanely euthanized and tissue specimens were collected for histopathological evaluation and immunohistochemistry (IHC), performed as previously described.^51^

## ACKNOWLEDGMENTS

This project is supported by TUBITAK-COVID Platform and Ministry of Industry, Science and Technology of Turkey.

